# Systematic identification of cancer driving signaling pathways based on mutual exclusivity of genomic alterations

**DOI:** 10.1101/009878

**Authors:** Özgün Babur, Mithat Gönen, Bülent Arman Aksoy, Nikolaus Schultz, Giovanni Ciriello, Chris Sander, Emek Demir

**Affiliations:** Computational Biology Center, Memorial Sloan Kettering Cancer Center, 1275 York Avenue, Box 460, New York, NY 10065, USA; Department of Epidemiology and Biostatistics, Memorial Sloan Kettering Cancer Center, 1275 York Avenue, New York, NY 10065, USA; Tri-Institutional Training Program in Computational Biology and Medicine, New York, USA

## Abstract

Recent cancer genome studies have identified numerous genomic alterations in cancer genomes. It is hypothesized that only a fraction of these genomic alterations drive the progression of cancer – often called driver mutations. Current sample sizes for cancer studies, often in the hundreds, are sufficient to detect pivotal drivers solely based on their high frequency of alterations. In cases where the alterations for a single function are distributed among multiple genes of a common pathway, however, single gene alteration frequencies might not be statistically significant. In such cases, we expect to observe that most samples are altered in only one of those alternative genes because additional alterations would not convey an additional selective advantage to the tumor. This leads to a mutual exclusion pattern of alterations, that can be exploited to identify these groups.

We developed a novel method for the identification of sets of mutually exclusive gene alterations in a signaling network. We scan the groups of genes with a common downstream effect, using a mutual exclusivity criterion that makes sure that each gene in the group significantly contributes to the mutual exclusivity pattern. We have tested the method on all available TCGA cancer genomics datasets, and detected multiple previously unreported alterations that show significant mutual exclusivity and are likely to be driver events.

---

Only a small fraction of genomic alterations present in a tumor are selected directly because of their ability to increase the cellular proliferation and to unlock barriers against growth and metastasis. The majority of the observed alterations, the so-called passengers, are indirectly selected due to incidental co-occurrance with a driver alteration or other selected event [1]. Differentiating drivers from passengers in cancer can help us identify tumorigenic mechanisms, drug targets, and design patient-specific therapeutic interventions.

Pivotal driver events, such as TP53 loss-of-function mutations, can be identified simply by their significantly high alteration rate in a set of tumors. More often, however, not one but several alternative driver alterations in different genes would lead to similar downstream events. In those cases the “selection bonus” is divided among the alteration frequencies of these genes. For current cancer genomics studies where the number of samples is two orders of magnitude smaller than the number of profiled genes per sample, statistical power of naive frequency based methods is not sufficient to differentiate these substitutive drivers from passengers. (Fig. 1).

**Figure 1:**
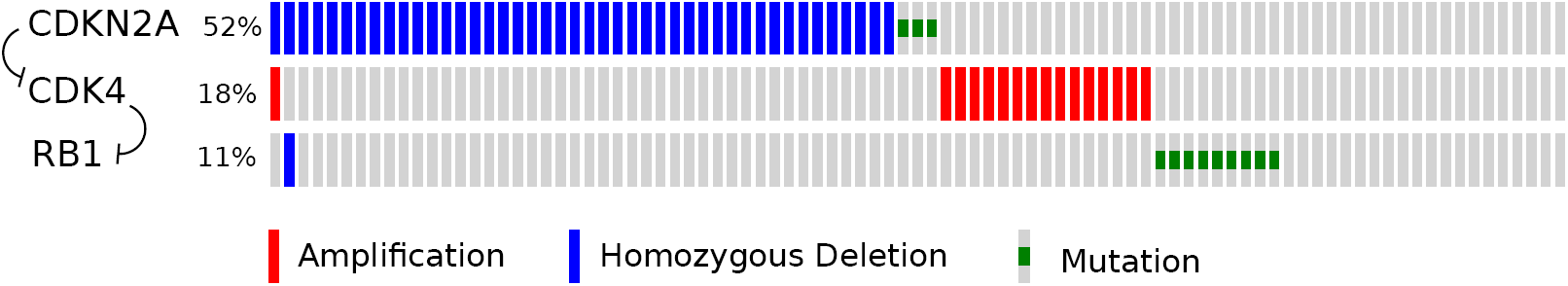
Distribution of CDKN2A, CDK4, and RB1 mutations and copy number changes in the TCGA Glioblastoma dataset [2], as provided by cBioPortal. At least one of the genes is altered in 78% of the cases, with overlap in only 2 samples. Even though RB1 is mutated only in 11% of the cases, its activity is potentially affected by alterations of the other two genes, which encode for upstream proteins in the signaling network.

A key observation is that when a member of a substitutive set is altered, the selection pressure on the other members is diminished or even nullified. As a result, we expect significantly less overlap in alterations of the alternative driver genes, creating a mutual exclusion pattern between their alterations. Supporting this expectation, it was previously shown that some functionally related genes are altered mututally exclusively in thyroid tumors [3, 4] and in leukemia [5].

This principle was first applied computationally by Yeang *et al.* to detect substitutive driver groups in cancer [6]. Their method calculates all pairwise mutual exclusion relations with a hypergeometric test. Miller *et al.* improved this approach by developing a statistical significance measure for the modules identified via pairwise exclusivity [7]. Ciriello *et al.* use the protein interaction graph for searching sets of mututally exclusive gene alterations [8]. They test each clique in the interaction network by random permutations to see if the observed overlap is significantly small. By using prior interaction knowledge, this approach can dramatically limit the search space. Vandin *et al.* suggest a weight function to score mutually exclusive alterations, which rewards coverage (number of samples altered in at least one of the genes in the group) while penalizing overlap [9]. They, then, search for subsets of genes that maximize the weight function. Zhao *et al.* and Leiserson *et al.* use the same weight function and expand on the search technique [10, 11]. Szczurek *et al.* propose a generative model for mutual exclusivity and test if the observed distribution of alterations fits this model better than a random model [12]. Their generative model assumes that genes in a module have equal chance to be altered, hence their result modules typically contain genes with similar alteration ratios.

We are expanding on these approaches by combining detailed prior pathway information with a novel statistical metric to improve both accuracy and biological interpretation and validation of the results. Specifically, we are using a large aggregated pathway model of human signaling processes to search groups of mutually exclusively altered genes that have a common downstream event. To enable this search, we also define a new statistical test that satisfies the following criteria:

i. Analytical: Scoring the mutual exclusion with random permutation testing is computationally expensive. Ciriello *et al.* can use such a metric for detecting significant cliques because the number of tested cliques is limited. For a wider search that includes non-clique subgraphs as in the case of upstream signaling pathways, the number of hypotheses that need to be tested increases by several orders of magnitude making the permutation testing infeasible.
ii. Soft: There are two kinds of mutual exclusivity defined in statistical literature: hard and soft. The hard mutual exclusivity [13] tests for two events that are assumed to be strictly mutually exclusive and the null hypothesis is that overlaps between them can be explained by random errors. Our assumed biological mechanism, however, implies a soft mutual exclusivity where two otherwise independent events overlap less than expected by random because of an interaction – in this case *partially* overlapping selective advantages. The soft mututal exclusivity of alterations of two genes can be statistically measured by a hypergeometric test, also known as Fisher’s exact test.
iii. N-ary: There is no consensus for testing the mutual exclusivity of more than two genes analytically. Yeang *et al.* test for all pairwise interactions to be significant in the group. This, however, is an overly strict test as a gene set can exhibit a strong mutual exclusion pattern as a group even if none of the pairs are significantly mutually exclusive. The weight function used in Vandin *et al.*, Zhao *et al.* and Leiserson *et al.* can test arbitrarily large sets. This metric, however, has a strong bias towards highly altered genes, and in some cases can select randomly occuring high coverage, high overlap sets, resulting in both false positives and negatives. We provide examples for these cases in the third section of the Supplementary Document.

Although our scoring metric can be applied to a wide spectrum of searches, we limit our scope to sub-graphs of molecular pathways where the members of the mutually exclusive group have a common downstream signaling target as defined in the public pathway databases. The rationale behind this is to focus on a subregion of the search space that has a higher density of true positives. At the cost of some recall, this reduction mitigates the statistical power loss due to multiple hypothesis testing. Another advantage of this reduction is that it nominates a “preliminary mechanistic explanation” for the observed mutual exclusivity - specifically a common effect on a downstream gene. It is, however, important to note that this is just one potential mechanism out of many–it should be treated as a starting point for futher biological inquiry. Statistical significance of the observed mutual exclusivity is independent from the hypothetical mechanism that nominated it for testing.

We tested our method on 17 different TCGA cancer datasets in cBioPortal [14], and identified multiple significant altered gene groups that are functionally related. We also present a comparison of performances of existing methods on simulated datasets. An implementation of the method in Java is freely available at http://code.google.com/p/mutex

## 1 Results

### 1.1 Measure for mutual exclusion

To measure mutual exclusion of a group of genes, we test each gene against the union of all other alterations in the group, and use the least significant p-value as the initial score of the group. To correct for multiple hypothesis testing, we estimate the member genes’ probability to have the observed p-value in a result group by chance, and derive the corrected p-values. We use the least significant of this second set of p-values as the final score of the group (Fig 2a, also see Online Methods).

**Figure 2:**
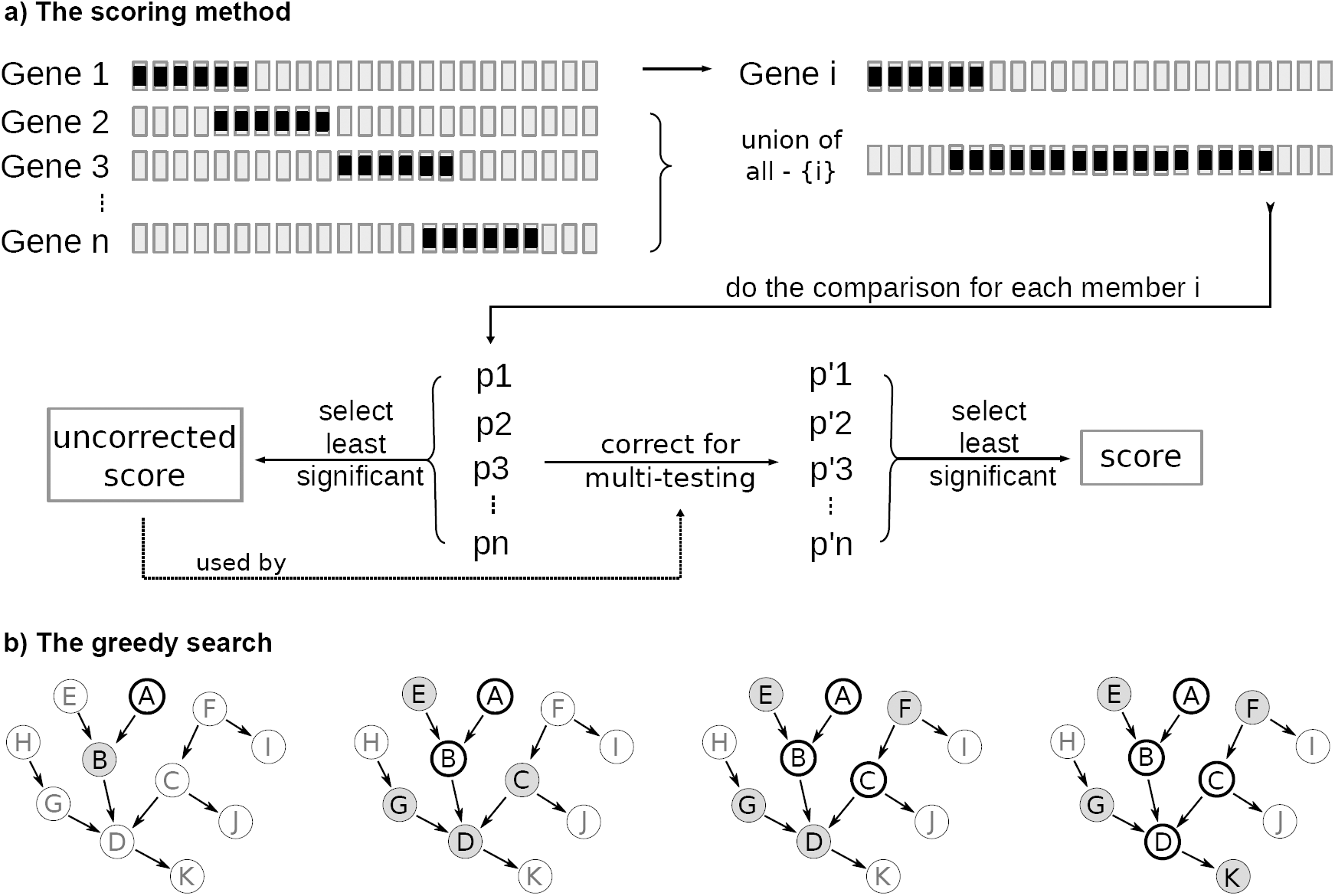
Searching and scoring method. (a) Mutal exclusivity of a group of gene alterations is evaluated by comparing each gene with the union of the other genes. The initial score is the least significant p-value. To correct for multiple hypothesis testing, we estimate the significance of the initial p-values during a search with permuted alterations. The least significant of these second p-values is the multiple hypothesis testing corrected group score. (b) At each step of the greedy search, we expand the group with the next best candidate gene from the surrounding genes that have a common downstream target with the group members, or they are a common downstream target themselves. In this illustration, 4 sample steps of the search are shown on a sample network. Thick-bordered genes are current group members and genes with gray background are candidates for the next expansion. The best scoring candidate gene is added to the group if it increases the score, and the candidates are re-assessed for the next phase. The search will stop if the group cannot expand anymore or a threshold group size is reached.

### 1.2 Searching the mutually exclusive group

We limit the search space for two reasons: testing every possible group of genes is not possible computationally, and even if we could, since the search space is combinatorially large, the significance of the results would be diluted due to the multiple hypothesis testing. Thus, we search for the mutual exclusivity only for groups where we have prior evidence for overlapping functions of genes. Our goal is not to predict new interactions but to find out which known pathways and or functions are altered in a certain cancer context and patients.

We built a large, directed gene network collecting interactions from Pathway Commons [15], SPIKE [16], and SignaLink [17] databases. This network is available in the project source code and its generation was described previously [18]. We use this network to search for groups of mututally exclusive genes that have a common downstream target on the network. We start the search by initializing a group with an altered gene as seed of the group, and greedily expand with the next best candidate gene. We define candidate genes such that after addition of a candidate, the members will still have a common downstream gene that can be reached without traversing any non-member genes (Figure 2b). Note that a common downstream gene can also be a member. We greedily expand the group with the candidate that best improves the group score. The search terminates when there are no remaining candidates or when the group size reaches a preset threshold. Algorithm returns a group and its score for each seed gene. To control the false discovery rate in the resulting groups, we estimate the null distribution of the final scores by running the same analysis on a set of permuted datasets, where gene alteration ratios and network connectivity are preserved, but sample distributions of alterations are shuffled.

We used this algorithm to identify mutual exclusion in the mutation and copy number profiles from 17 TCGA studies [2, 19, 20, 21, 22, 23, 24, 25], deposited in cBio Portal [14]. We cropped the gene network to the proximity of significantly mutated genes (provided by MutSig [26]) and copy number significantly altered genes (provided by Gistic [27]). Here, proximity means the neighbor genes and the genes that have a common downstream target. Thresholds 2 and −2 were used for copy number amplification and deletion, respectively, for discretized Gistic values. We only used the copy number changes which are confirmed by an expression change (see Online Methods). For each study, we filtered out genes that have a low alteration rate to reduce the noise in the data. We searched groups up to size 5. We ran 10000 permutations for estimating the null distribution of the member p-values of genes in groups, and we used 100 iterations for estimating the null distribution of the final group scores in the result. For each study, we selected the false discovery rate cutoff that maximizes the expected value of *true positives – false positives* in the results.

### 1.3 Results on TCGA datasets

We provide the analysis results of individual studies in the Supplementary Document. The distribution of alterations in endometrial cancer samples is exceptional in the sense that samples are strongly dominated by either copy number alterations, or mutations (Suppl. Fig. 1). Because of this, many of the copy number changes are mutually exclusive with many of the mutations. Since mutual exclusion of the copy number alterations and mutations result from a higher order event, the mutual exclusivity for these two subtypes have little additonal biological implication. To remove the effect of subtypes, we divided the endometrial cancer samples into two, and treated them as different studies.

We observe that 31 genes appear in the results of at least two studies. Figure 3 shows these recurrent genes and their recurrent co-presence in the same result group. Not surprisingly, the most recurrent gene in the results is TP53, and it is followed by PTEN, KRAS, MYC, PIK3CA, BRAF, EGFR, and NRAS–all well-known tumor suppressors and oncogenes. The next most recurrently found two genes are OBSCN and ARID1A. OBSCN functions in myofibrillogenesis, and known to activate Rho GTPases [28, 29]. Motif-based studies also predict it to bind to PIK3R1–the regulatory component of PI3K complex [30]. ARID1A was previously shown to be a tumor suppressor in gastrological cancers. Two other genes that are relatively less associated with cancer are LAMA2 and SPTB. LAMA2, the alpha subunit of laminin, functions in cell attachment and mobility. It is also known to function in a complex that activates Rho GTPases [31]. SPTB is a member of the spectrin family, which are membrane cytoskeletal proteins that functions in cell membrane organization and stability. It was previously shown that another spectrin, SPTBN1, functions in TGF-beta signaling, and its loss can contribute to hepatocellular cancer [32].

**Figure 3:**
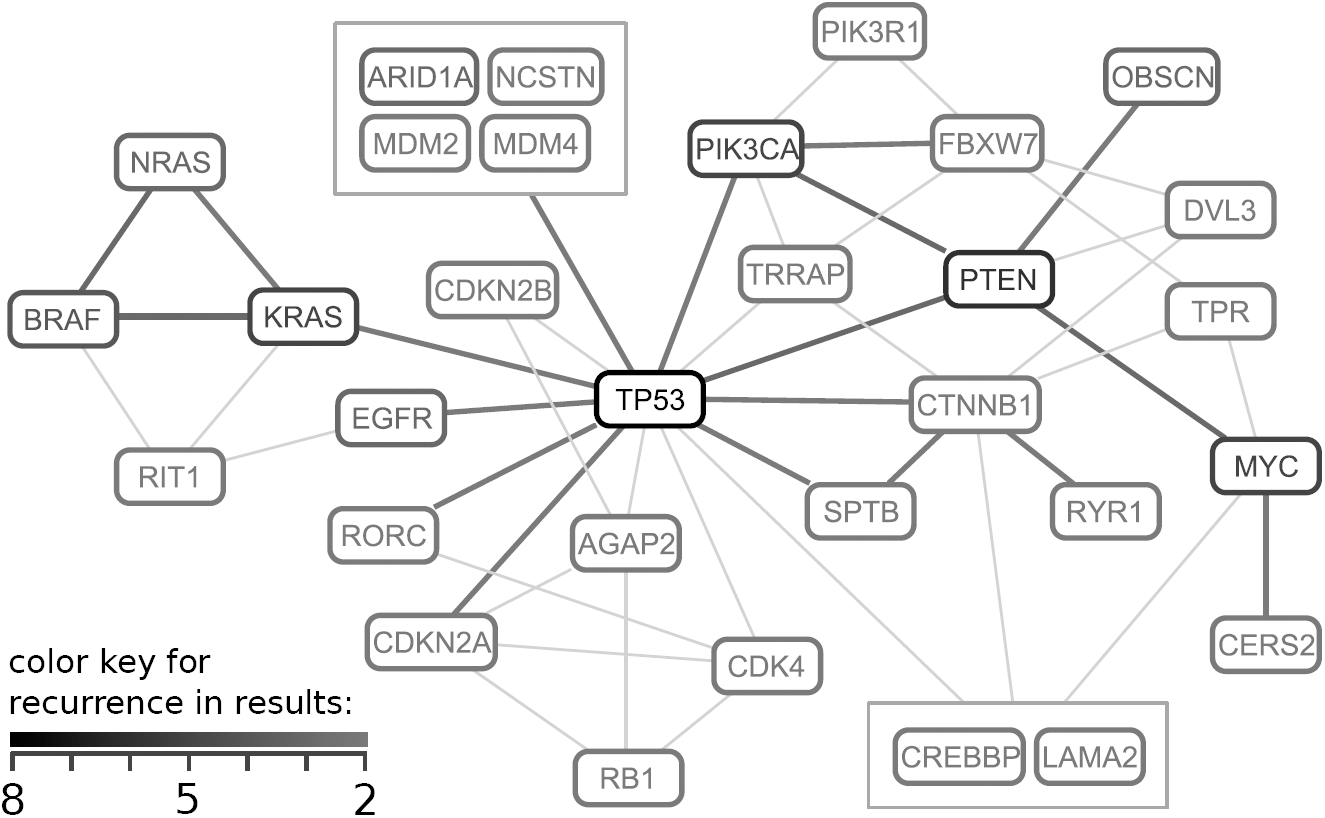
A network constructed using recurrent genes in the results. The genes in the graph appears in results of at least two different studies. Thick edges represent recurrent co-presence of gene pairs in the same mututally exlusive set. Thin edges represent non-recurrent co-presence of gene pairs, and only used to connect the genes that lack a recurrent edge. Note that there are many other non-recurrent edges between recurrent genes, omitted in the graph for reducing complexity. See the Suppl. Fig. 44 for a complete graph with all non-recurrent genes and co-presences. Figure prepared using ChiBE [33].

The most frequent common targets in the result groups are PIK3R1, HRAS, BRAF, MYC, RAC1, and RHOC. Figure 4 shows 5 result groups that are at the upstream of RHOC. RHOC is a member of Rho GTPases, whose members function in the regulation of the cell shape, attachment and motility. Even though RHOC alterations are not frequent in TCGA samples, its over-expression was previously shown to promote metastasis of cancer cells [34, 35]. We observe that it is expressed in great majority of TCGA samples (Suppl. Fig. 45). These mutual exclusive alterations at the signaling upstream of RHOC in several different cancers suggest that activation of RHOC can be one of the major downstream effect of driver alterations.

**Figure 4:**
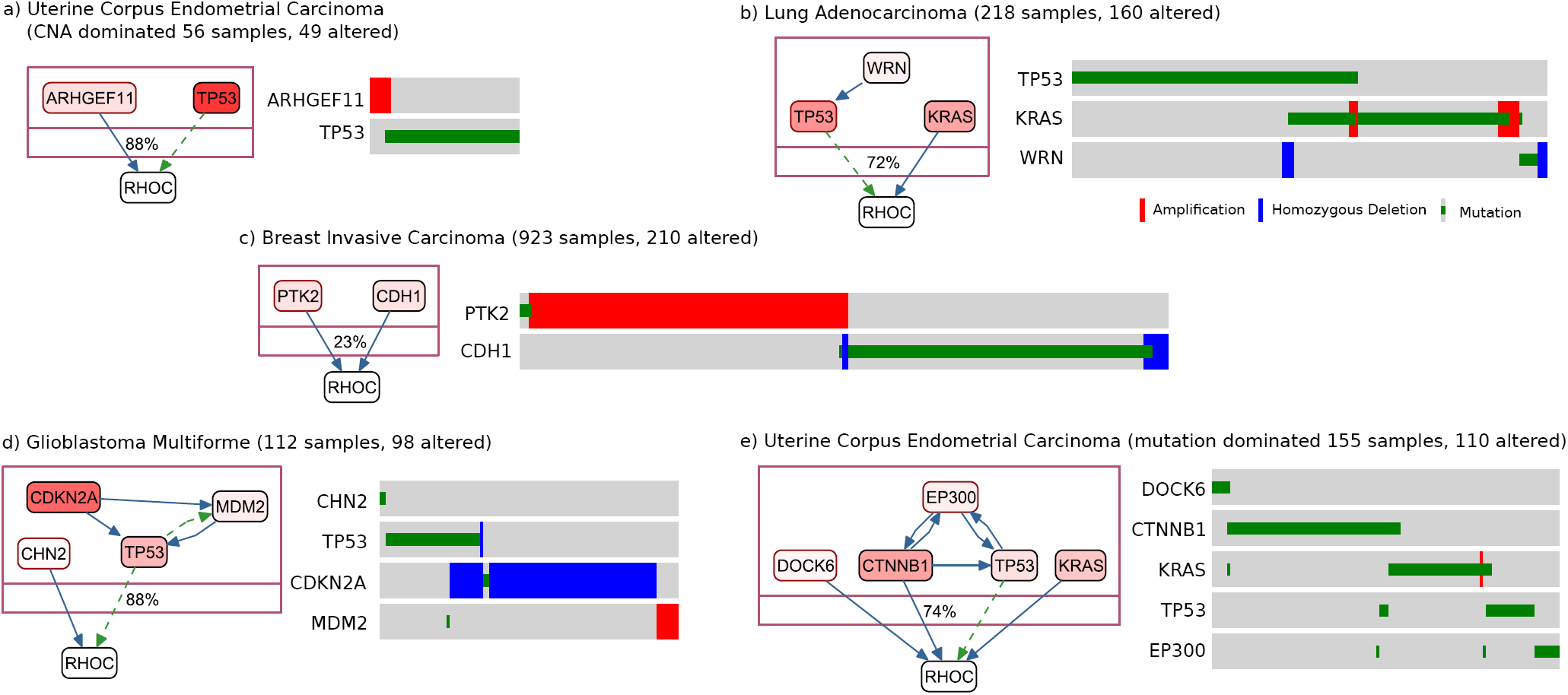
Sample result groups that have RHOC as a common downstream target. Oncoprints are shown at the right of each group, where unaltered samples are ommitted. Gene color intensities are proportional to gene alteration ratios. On the signaling network, dashed green edges represent transcriptional relations, and solid blue edges represent post-translational relations. Genes in the result groups are shown inside a compound node, whose label shows the alteration-coverage of samples in the group. This is also equal to the visible portion of samples on the oncoprints.

### 1.4 Comparison of methods that detect mutual exclusive gene alterations

We compared the performance of our method (Mutex) with performances of previously published methods – Pairwise search [6] (2008), RME [7] (2011), Dendrix [9] (2012), MEMo [8] (2012), MDPFinder [10] (2012), Multi-dendrix [11]
(2013), and ME [12] (2014) – on simulated datasets.

In our first trial, we derived a large dataset from the breast cancer dataset in cBioPortal (using mutations and expression-confirmed copy number changes), which contains 830 genes with an alteration rate of at least 3% in 958 samples. The derivation steps are: (i) randomize sample distribution of gene alterations while preserving alteration ratios, (ii) determine 50 non-overlapping groups of genes, each with 3 members that are upstream of a common target in the signaling network, (iii) iterate over gene alterations in groups and re-randomize the overlapping alterations, and repeat that once more. The last step decreases the chance of overlaps in the seeded groups. With this large data set, we were not able to obtain results from MEMo and ME methods – both algorithms did not scale to the large data size. A comparison of ROC curves (Figure 5) shows a dramatic improvement over existing methods.

**Figure 5:**
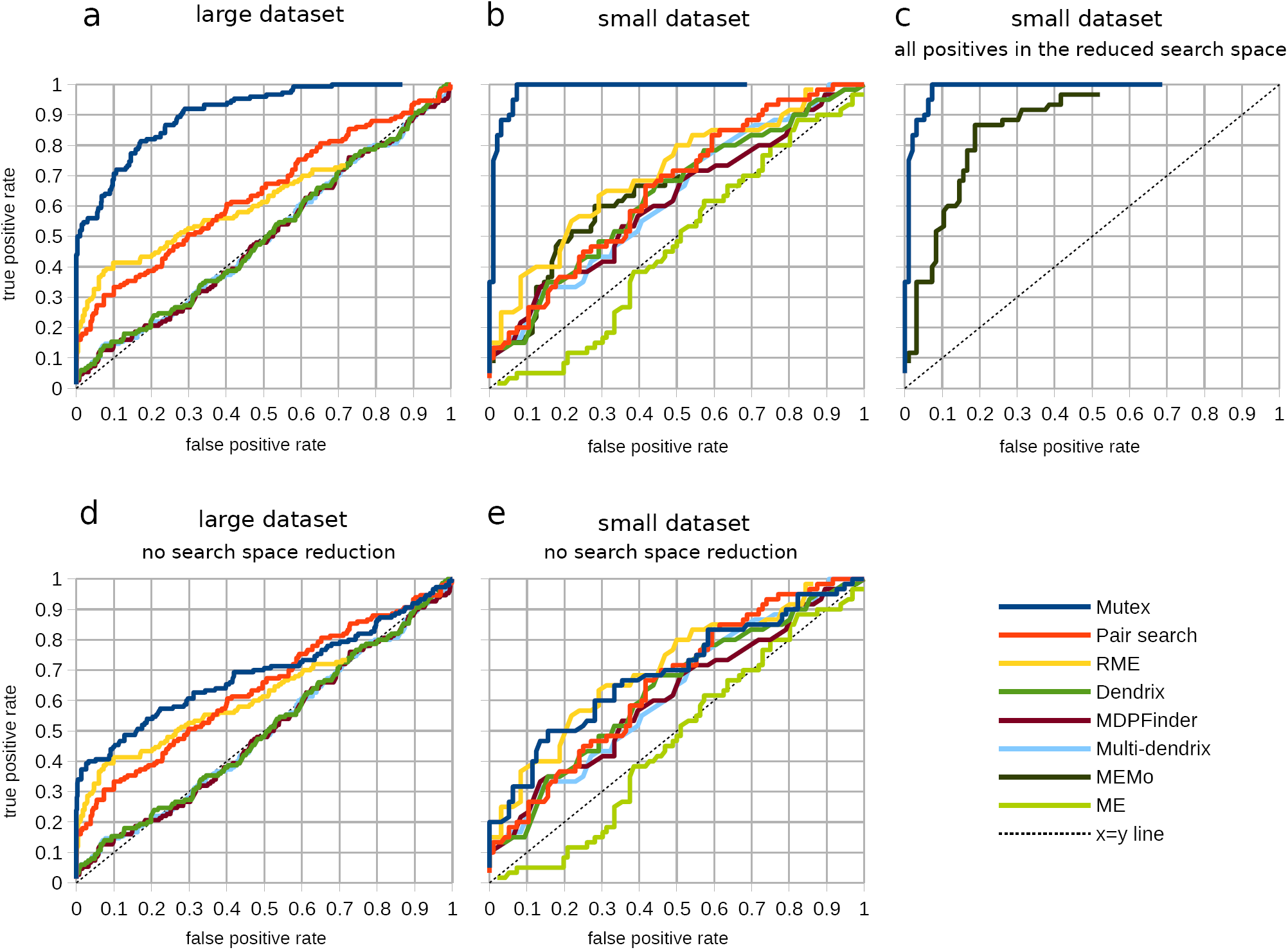
ROC curve comparisons of methods that detect mutual exclusive alterations on simulated datasets. a) Comparison on the large dataset. b) Comparison on the small dataset. c) Comparison of methods that do search space reduction (Mutex and MEMo) after ensuring all seeded groups to be in the reduced search space of both methods. d,e) Comparisons after removing the search space reduction of Mutex, on large and small datasets.

Mutex outperforms other methods due to two reasons: (i) reduction in search space, (ii) a more precise search metric. In order to understand how these two components contribute to the improvement we compared a modified version of Mutex that does not use the signaling network and does not reduce the search space, with other methods that do not reduce search space (Fig. 5d,e). As expected, the ROC performance of Mutex decreased and became comparable with the other methods. The difference, however, is still substantial in terms of the number of results in a realistic use case. If we use a 5% FDR cutoff on the large dataset, modified Mutex recovers 49 true positive genes in the results while next-best method, RME recovers only 12.

This experiment validates that Mutex’s statistical metric is an improvement over other methods even without search space reduction. It also clearly demonstrates that exploiting the prior pathway information can improve precision. There is one caveat though – in real data, common downstream molecule is one possible pattern out of many that can lead to mutually exclusive alteration patterns. The algorithm, however, is pattern agnostic and can be readily extended to other patterns – a research direction that we are exploring.

In order to include MEMo and ME to the comparison, we prepared a smaller simulated dataset (156 genes and 463 samples), and seeded 20 groups, each with 3 members, using the same procedure, and compared ROC curves (Fig. 5b). Again, Mutex outperformed all other methods. In these comparisons, most of the advantage of Mutex is coming from reducing the search space. MEMo is the only other method that reduces the search space. MEMo, however, has a disadvantage in this comparison because we did not select the seeded groups to also appear as nodes of a clique in the interaction network that MEMo uses. For a fair comparison, we added new interactions between seeded group members to MEMo’s interaction network database. This modification creates a clique for each seeded group that can be detected by MEMo. In this case (Figure 5c), the performance difference between Mutex and MEMo is mostly due to the more stringent metric used by Mutex that ensures each member contributes significantly to the group thus decreasing the number of false positives.

Performance tests also demonstrates that Mutex scales very well to large datasets and large groups, both in terms of memory usage and runtime. It has similar runtime characteristics with Multi-dendrix and MDPFinder and is much more efficient than RME, Dendrix, MEMo and ME.

## 2 Discussion

We have developed a method that can detect mutually exclusive genomic alteration patterns in cancer genomic datasets. Mutex is unique in its ability to use prior pathway information efficiently to search the graph structure and reduce the search space. This reduction trades recall for precision. Given the highly noisy nature of the cancer genomic datasets, this is almost always a desirable trade-off. We also offer a new efficient and better statistical test that, even without the prior pathway information, improves the existing approaches.

Our approach detects many interesting genes and groups, that are candidate cancer drivers, that would not be detected by frequency based methods such as MutSig. Additionaly, mutex groups couples less known, less frequently altered genes with well characterized cancer drivers, suggesting a mechanism of action. We also observe that many result groups are overlapping. We speculate that this suggests a highly coupled selection advantage between genomic modifications as opposed to well defined modules or pathways.

Our mutual exclusivity score for groups of genes is analytical and fast to calculate for a single group. We still use permutation testing for multiple hypothesis correction when testing multiple groups. Since our method uses the same estimated null distribution for a gene regardless of the tested group, however, our approach scales substantially better compared to approaches where a permutation testing is performed for each evaluated group.

We plan to extend this work towards searching other topological structures on the biological network. The current method selects genes with a common downstream target, and requires all group members to be directly linked on the network without a non-member linker node. Allowing linker nodes can help identify more distant mutual exclusion relations.

## 3 Methods

### 3.1 Mutual exclusivity of a pair

We define alteration of two genes to be mutually exclusive if their overlap in samples is significantly less than expected by random (Suppl. Fig. 46). The statistical significance of the low-overlap can be calculated using a hypergeometric test.

For a pair of genes, a hypergeometric test is an analytical alternative to permutation testing, where permutations remove the dependency between alterations while preserving alteration rates, and assume equal probability for each sample to be altered. This assumes a uniform alteration frequency from sample to sample. This might not always be the case, especially for the so-called “hyper-mutated” samples, often caused by a preceding mutation in DNA repair mechanisms. Properly addressing this heterogeneity is very complicated, as one should account for the fact that each overlap has different probability in the null model. This is still an open problem. At the cost of statistical power, we partially mitigate this issue by excluding hyper-altered samples from the analysis. Specifically, we removed samples that have more alterations than Q3 + (1.5 x IQR) where Q3 is the third quartile in the distribution and IQR is the interquartile range. This criteria is often used in box plots to mark outlier values.

### 3.2 Mutual exclusivity of a group

There is no standard way of testing whether a group of more than two genes exhibits a mutual exclusion pattern. We can compare pairs of genes, or we can get the union of alterations of a subset of genes in the group and compare to another disjoint subset. There are combinatorially many ways to test the significance of the mututal exclusion in a group of genes. Among a wide number of possibilities, here we identify a subset of these tests to measure mutual exclusivity.

To make sure that every member of the group significantly contributes to the pattern, we use the following null hypothesis:

*H*_0_ : *The specific member gene in the group is altered independently from the union of other alterations in the group.*

We test *H*_0_ for each member, by evaluating the co-distribution of the gene with the union of other genes (Figure 2a) using a hypergeometric test. For a group of *n* genes, this method generates *n* p-values, which are probabilities for independent distribution of each member gene. Since we would like each member to contribute to the pattern, we use the least significant member p-value as the initial score of the group. A closed-form expression for this metric is provided in Equation 1, where *g*_*i*_ is the alterations of *i*^*th*^ gene in the group, *g*_*n−i*_ is the merged alterations of group members excluding the *i*^*th*^ gene, and *H* is the hypergeometric test that generates the p-value of mutual exclusivity of two array of alterations.

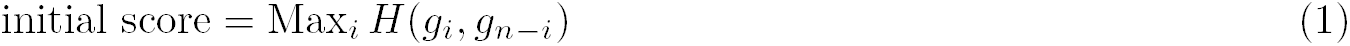

Since we are testing more than one group, this initial (uncorrected) score is affected by multiple hypothesis testing. To correct for multiple hypothesis testing, we first estimate the null distribution of the initial p-values of each gene, then calculate the significance of the observed initial p-value for each member. Among this second set of p-values, we select the least significant as the multiple hypothesis testing corrected final score (Figure 2a).

To estimate the null distribution of the initial p-value for a gene, we sample it by permuting alterations of that gene, and searching for the group with the maximum uncorrected score using the same greedy search method. We only permute the alterations of the gene in question, because we are testing that gene in its specific network environment with a specific alteration pattern in surrounding genes.

To be able to control the false discovery rate (FDR) of the resulting groups, we need a measure for significance of the final score. Most popular FDR methods, like the Benjamini-Hochberg procedure, are developed for p-values that are assumed to have a uniform null distribution. The final scores do not have a uniform null distribution even though they were derived from p-values. This is both due to selecting the least significant p-value in the group (which shifts the null distribution to the right) and searching for the best scoring set of the seed gene (which shifts the null distribution to the left). Since it is hard to estimate the shape of the null distribution analytically, we estimate it by running the analysis with all-permuted gene alterations, multiple times. Using the estimated null distribution of the final scores, we select the most significant results to satisfy a certain false discovery rate.

### 3.3 Verifying copy number alteration data with expression

To reduce the noise in the data, we filter out the copy number changes that are not supported by gene expression. We compare gene expression of copy number intact samples with samples that have the specific copy number alteration (either amplification or deletion) (Supplementary Fig. 47). If the difference of mean expressions between two distributions is not in the expected direction or it is not statistically significantly different using a t-test with a 0.05 p-value threshold, we do not use the copy number alterations in question. If the difference between the means is significant, we determine the expression threshold where expression is more likely to belong to the copy number changed distribution, and use only the copy number changes with an expression satisfying the threshold.

### 3.4 Filtering genes

We filtered out genes that are not at the proximity of recurrently mutated or recurrently copy number altered genes on the signaling network. We used the MutSig analysis results with q-value threshold 0.05 for recurrent mutations, and used Gistic results with q-value threshold 0.05 for recurrent copy number alterations, both obtained through Broad Firehose. We define proximity genes as the union of the neighbor genes and the genes that have a common downstream target (the upstream of the downstream genes). We used following procedure.

*A* = *MutSig genes*

*B* = *Gistic genes*

*C* = *Proximity*(*A*)

*D* = *B* ∩ *C*

*E* = *A* ∪ *D*

*F* = *Proximity*(*E*)

*Result* = *E* ∪ *F*

For performance purposes, we filtered out genes with very low alteration rates. We used a minimum threshold of 0.01. The threshold was increased for studies that have very high overall alteration frequencies to ensure that number of seed genes remains less than 500.

### 3.5 Generation of simulated datasets

We derived the simulated datasets using the two TCGA breast cancer datasets deposited in cBioPortal. For the larger simulated dataset, we used the dataset named “Breast Invasive Carcinoma (TCGA, Provisional)”, which contained 958 complete samples at the time. Here, complete means that the sample have mutation profile, copy number profile, and expression profile available. We used mutations and expression-verified copy number alterations as previously described. We filtered out genes with less than 3% alteration rate, which gave us 830 altered genes. For the small dataset we used the dataset named “Breast Invasive Carcinoma (TCGA, Nature 2012)", which contains 463 complete samples. We filtered out genes with less than 5% alteration, which gave us 158 altered genes.

We randomized the alterations of each gene separately, preserving the number of alterations, but re-assigning sample distributions. We chose 50 non-overlapping positive groups for the large dataset, and 20 positive groups for the small dataset, which are composed of 3 genes that have a common downstream gene on the signaling network. We applied the below algorithm two times to artificially introduce mututal exlusion to the alterations of positive groups by moving the overlapping alterations to new random locations.

~~~
**for** each positive group G **do**
 **for** each member gi of G **do**
               **for** each member *g*_*j*_ of G where *i* ≠ *j* **do**
                        **for** each sample k **do**
                               **if** both *g*_*i*_ and *g*_*i*_ are altered in sample k **then**
                                     *x* ← random sample where gi is not altered
                                     alter gi at sample x
                                        un-alter gi at sample k
                                **end if**
                          **end for**
                **end for**
     **end for**
**end for**
~~~

## Acknowledgements

This research was supported by NIH grants (U41HG006623) and (GM103504).

## Contributions

ÖB and ED conceived the idea. ÖB, ED and MG developed the method. ÖB implemented the method with help from BAA. CS, NS, BAA, and GC contributed to the discussions during the course of the project.
The manuscript was written by ÖB, ED, and NS.

## Competing interests

The authors declare that they have no competing financial interests.

